# Microsaccades and attention in a high-acuity visual alignment task

**DOI:** 10.1101/592709

**Authors:** Rakesh Nanjappa, Robert M. McPeek

## Abstract

While aiming and shooting, we make tiny eye movements called microsaccades that shift gaze between task-relevant objects within a small region. However, in the brief period before pressing trigger, microsaccades are suppressed. This might be due to the lack of the requirement to shift gaze as the retinal images of the two objects start overlapping on fovea. Or we might be actively suppressing microsaccades to prevent any disturbances in visual perception caused by microsaccades around the time of their occurrence and their subsequent effect on shooting performance.

In this study we looked at microsaccade rate while participants performed a simulated shooting task under two conditions: normal viewing in which they moved their eyes freely and eccentric condition in which they maintained gaze on a fixed target while performing shooting task at 5° eccentricity. As expected, microsaccade rate dropped at the end of the task in the normal viewing condition. However, we found the same for the eccentric condition in which microsaccade did not shift gaze between the task objects.

Microsaccades are also produced in response to shifts in covert attention. To test whether disengagement of covert attention from eccentric shooting location caused the drop in microsaccade rate, we monitored participant’s spatial attention location by employing a RSVP task simultaneously at a location opposite to the shooting task. Target letter detection at RSVP location did not improve during the drop in microsaccade rate, suggesting that covert attention was maintained at the shooting task location.

We conclude that in addition to their usual gaze-shifting function, microsaccades during fine acuity tasks might be modulated by cognitive processes other than spatial attention.

## INTRODUCTION

We use rapid eye movements known as saccades to shift our gaze serially between multiple regions of interest (ROIs) in our visual field, which then guides subsequent motor behaviors like navigating, or reaching and grasping objects. The amplitude of these saccades during viewing of a particular scene depends on the separation between ROIs in that scene. In natural scenes, ROIs are widely spread out, and thus we typically make saccades that are 4° or larger (Dorr, Martinetz, Gegenfurtner, & Barth, 2010). However, in some tasks that require high visual acuity, like threading a needle or aiming a rifle, ROIs may be separated by distances of less than 1°. In such tasks, we use saccades as small as 20 minutes of arc to shift gaze precisely and to explore a narrow range of space. These small saccades can then be used to guide fine motor adjustments, just as larger saccades do (Ko, Poletti, & Rucci, 2010). Saccades falling in this small range, known as microsaccades, enjoy a special status in the field of eye movements for reasons different from their exploratory nature described above: microsaccades are also produced at a rate of 1-2 per second while trying to hold gaze on a fixation target. The possible function of these fixation saccades has been a matter of debate (e.g., Collewijn & Kowler, 2008; Rolfs, 2009). In the contexts of both exploration and fixation, modulations in the spatiotemporal properties of microsaccades have been shown to reflect different phenomena. Although changes in microsaccade rate and direction during fixation reflect shifts of covert attention (Hafed & Clark, 2002; Engbert & Kliegl, 2003a), their most obvious function, that of relocating gaze, is uncovered only in high visual acuity tasks. When it is necessary to precisely explore a narrow region of space, such as when threading a needle, both the average and the instantaneous microsaccade rates are suppressed (Winterson & Collewun, 1976; Bridgeman & Palca, 1980; Ko et al., 2010). This observation has led to opposing interpretations regarding the effect of microsaccades on task performance, and their role in general.

Winterson & Collewijn (1976) recorded the eye movements of human subjects while they performed two separate fine acuity visuo-motor tasks: threading a needle and shooting a rifle. They made two important observations in both tasks: first, average microsaccade rate during these tasks was lower than during prolonged fixation on a fixation target. Second, within the time course of a trial, microsaccade rate decreased with time, with almost no microsaccades made in the final second of the task, i.e., just before subjects inserted the thread in the eye of the needle or pressed the rifle trigger. Based on these observations, they concluded that microsaccades are detrimental to performance in tasks requiring high visual acuity and are thus suppressed. Similar conclusions were drawn by another study which asked subjects to passively view the motion of a needle and thread (without any motor control), and to make a perceptual judgment about their alignment (Bridgeman & Palca, 1980). Thirty years later, Ko et al. (2010) designed a simulated version of the needle-and-thread task in which subjects freely viewed the task stimulus on a monitor and controlled the vertical position of a thread approaching a fixed needle at a constant horizontal velocity. They made the same two observations regarding microsaccade rate, but drew different conclusions. First, they suggested that microsaccades produced during attempted fixation served a different purpose than those produced during the needle-and-thread task, and hence their comparison cannot be used to draw any conclusions. Second, through a detailed spatial analysis of the microsaccades produced during an earlier period in the trials, they showed that microsaccades precisely relocated gaze according to the temporally changing separation between the ROIs, and thus served the dynamic needs of gaze relocation over a very narrow region. Based on this, they hypothesized that microsaccade rates dropped at the end of the trial not because they were detrimental to the task, but because at that point, both ROIs overlapped on the effective foveal region, thus obviating the need for any further gaze shifts.

In our present study, we simulated a shooting task in which subjects controlled the motion of a gun sight so as to align its center with the center of a stationary shooting target. To study the effects of microsaccades’ gaze-relocating function on their rate, we dissociated the gaze-relocating function of microsaccades from their occurrence by asking subjects to perform the same task in two different viewing conditions. In the normal viewing condition, as usual, microsaccades shifted gaze according to the ongoing demands of the task, and their rate dropped at the end of the trial, as reported in earlier studies. In an eccentric viewing condition, subjects maintained fixation on a central fixation target while the shooting task stimuli were presented at a 5° eccentric location. As a result, subjects used peripheral vision to view the task stimulus, and thus, any microsaccades produced during the task could not serve the purpose of relocating the fovea between the peripherally-viewed ROIs. Nevertheless, we observed a similar drop in microsaccade rate. This suggests that there is something other than a gaze-relocation demand which suppresses microsaccades during the end of the eccentric viewing task. We speculated that this decrease in saccades in the eccentric viewing task may reflect the disengagement of attention from the peripheral shooting task stimuli. However, in a final experiment, we tested this explanation, and found that the drop in microsaccade rate in the eccentric viewing condition does not appear to reflect a release of attentional disengagement from the peripherally-attended task location. Put together, our findings suggest that microsaccade production in such tasks is affected by factors other than just their gaze-relocation function, and that the exact cause of their suppression remains a topic for future research.

## METHODS

### Participants

Seven (4 female) students from the Graduate Center for Vision Research, SUNY College of Optometry, with normal or corrected to normal vision and no known oculomotor defects, ranging in age from 25-30 years, participated in Experiment 1. Five of these subjects also participated in Experiment 2. Each participant signed a consent form approved by the SUNY College of Optometry Institutional Review Board. Participants received a base payment of $10 per experimental session plus an additional amount contingent upon their performance in the task, with the total payment not exceeding $20 for a single session. Although participants were not totally naïve about the purpose of the study, they did not have any prior experience of participating in a similar task or one which could have altered their microsaccade strategy in a fine acuity visuomotor task.

### Task

Participants sat 120 cm away from an IPS LCD monitor (Cambridge Research Systems Display++; 71 × 39.5 cm, 1920 × 1080 pixels, refresh rate 120 Hz, gray background) in a room with ambient lighting. Their heads were stabilized with a chin and forehead rest, and monocular eye movements (left eye for all participants) were recorded using an EyeLink 1000 (SR Research) at 1000Hz.

#### Experiment 1

Participants performed a simulated shooting task under two viewing conditions, each in a separate session consisting of 120 trials and lasting for ∼40 minutes, conducted on separate days. Each session began with a practice block in which participants performed 20 trials to become familiarized with the task and the associated push-button controls. After the practice block, the eye tracker was calibrated using Eyelink’s standard 9-point calibration. In the first condition, viz., the normal viewing condition, each trial started with the presentation of a 10’ wide black circular fixation target at the center of the screen (**Fig. 1**). Participants maintained fixation on the fixation target for a duration of 5 seconds, during which they were instructed not to blink. A blink resulted in termination of the trial, and a fresh trial began. After this prolonged fixation period, the fixation target disappeared and the shooting target (a black disc subtending 10’) appeared at one of 5 possible positions (0°, 5° left/right, 3° up/down), along with the ‘sight’ (a black circle outline of diameter 1° and boundary width 1.4’) within close vicinity of the shooting target. The starting position of the sight was randomly picked from an invisible square boundary of length 1.5° centered on the shooting target. Immediately after presentation, the sight started to move in a randomly-selected diagonal direction (45°, 135°, 225°, or 315° in direction with respect to the target) with a fixed velocity (randomly selected to be 9, 12, 13 or 15 min/sec). Participants used 4 direction buttons on a gamepad to control the sight’s direction of motion (which was constrained to the four directions listed above). The goal was to align the center of the moving sight with the center of the fixed shooting target. A single button press resulted in a corresponding change of direction (e.g., pressing the left button set the horizontal component of the motion to the left) with no change in overall speed. Participants pressed a ‘shoot’ button on the gamepad when they judged the centers of the two objects to be perfectly aligned. Following this ‘shoot’ event, the task stimuli disappeared, and performance on the trial was reported at the location of the fixation point as a score (out of 10) calculated based on the distance of the sight center from that of the target. Task eccentricity and sight velocity for each trial were picked randomly, and with equal probability, from the discrete values listed above.

**Figure 1.**
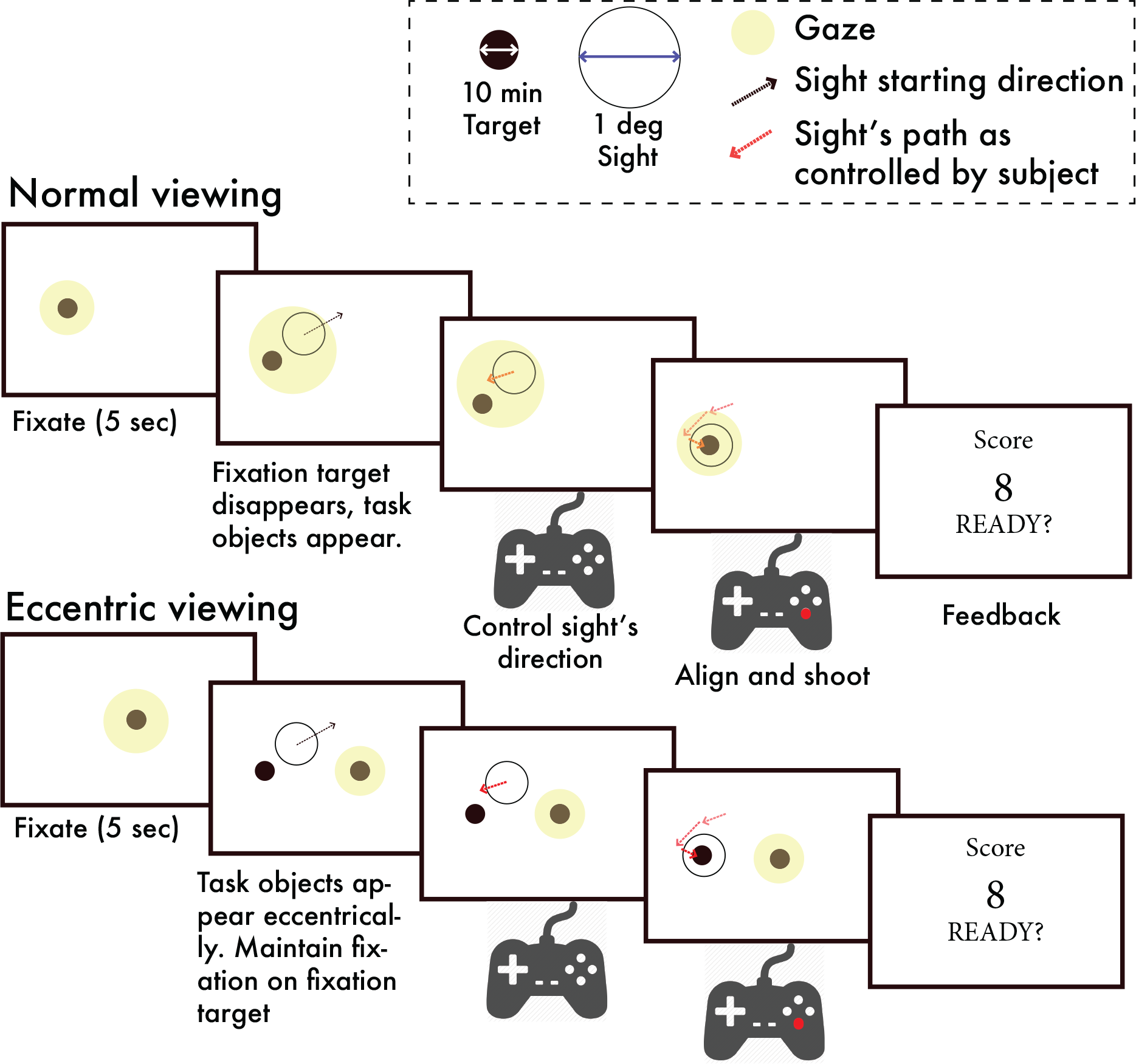
Experiment design. In Experiment 1, during the normal viewing condition, a trial started with fixation on a central fixation target for 5 sec, followed by presentation of shooting task stimuli, during which participants were free to move their eyes, and adjusted the sight’s direction of motion using a gamepad. The yellow patch shows the likely gaze position of the subject. The black arrow shows the initial motion path of the sight and the red arrows shows the motion path after a participant adjusted its direction. After aligning the sight’s center to the center of the shooting target they pressed the ‘shoot’ button on the gamepad to end the trial, following which their performance in that trial was reported to them as a score out of 10. In the eccentric viewing condition, everything was similar except that shooting task stimuli always appeared at an eccentric location and participants were required to maintain their fixation on the central fixation target while viewing the shooting task stimuli using peripheral vision. Score was presented at the fixation location.

In the second condition, viz., the eccentric viewing condition, a trial started with a 1-second fixation period, after which the shooting target and sight appeared at one of the four eccentric task locations (5° left/right, 3° up/down), while the fixation target remained on the screen. As opposed to the previous condition, in which participants were free to move their eyes, subjects were now instructed to maintain fixation on the central fixation target, and to use their peripheral vision to accomplish the same task (i.e., to align the sight center with the target center and shoot). Fixation was monitored using a fixation-check window, which consisted of an invisible square boundary of length 2° centered on the fixation target. A trial was aborted if the eye moved out of this window, or if the participant blinked. Participants could take a maximum of 30 seconds to finish a trial, and they controlled the beginning of the next trial with a button press. Calibration was re-done between trials if subjects moved their heads significantly. A session ended with completion of 120 valid trials.

#### Experiment 2

This experiment consisted of a dual-task situation, in which participants performed the same simulated shooting task as in Experiment 1, while simultaneously performing a task that required detecting a target letter from a stream of rapidly and serially presented letters (Rapid Serial Visual Presentation [RSVP] task). The RSVP stream consisted of 1° wide letters of the English alphabet, presented at a frequency of 5 Hz. The letter stream for each trial consisted of a random sequence of non-target letters, with the target letter interspersed such that target letter frequency was 0.5 or 0.8 Hz. The target letter remained the same for a given session. Participants reported detection of a target letter by pressing a button on the gamepad as soon as they saw it.

They performed this dual task under two viewing conditions with respect to viewing of the shooting stimuli. In the normal viewing condition (50% of trials), the shooting stimuli appeared at the center of the screen, where subjects were fixated, and the RSVP task stimuli appeared at one of two eccentric locations (5° left/right), and were viewed peripherally. In the eccentric viewing condition, participants maintained fixation on the central fixation target while the shooting and RSVP task stimuli appeared at eccentric locations on opposite sides of fixation. Again, fixation was monitored used a fixation-check square window of length 4° centered at the central fixation location, and a trial was aborted if participants blinked or moved their eyes out of this boundary. Participants adjusted the sight’s direction of motion as before, using the 4 direction buttons, and reported RSVP target detection using another button. Finally, they pressed the ‘shoot’ button when they judged the sight and target to be aligned. Following this, both task stimuli disappeared, but instead of the trial ending immediately, an extra 1 second was made available to report any last-moment RSVP target detection made by subjects. Following this, their shooting performance and RSVP task performance (percentage target letter detection) were reported to them at the location of fixation.

## RESULTS

In a simulated shooting task, participants used a gamepad to align the center of a moving circle (hereafter referred to as the ‘sight,’ as in the gun sight through which shooters view the shooting target to take an aim) with that of a fixed target disc (referred to as the shooting target). A trial started with the sight moving diagonally in a random direction with a constant velocity, with an initial separation of 1° between the target and sight (two regions of interest [ROIs]). Participants adjusted the sight’s direction of motion so as to align its center with that of the target, and then ‘shot’ by pressing a ‘shoot’ button at a time when they perceived the centers of the two objects to be perfectly aligned. They performed this task under two conditions of viewing: (i) normal viewing, in which participants were free to move their eyes anywhere on the screen, and (ii) eccentric viewing, in which they maintained fixation on a central fixation target and viewed the task stimuli with their peripheral vision. In the normal viewing condition, each trial was preceded by a 5 second long fixation period on the central fixation target which was used as a control condition.

Participants took a longer time before shooting, but performed better when they viewed the task stimuli normally as compared to when they used their peripheral vision. In the normal viewing condition, they finished a trial in 5.93 ± 1.28 sec (mean ± s.d.) whereas they took only 4.53 ± 0.23 sec in the eccentric viewing condition (paired t-test, t(6) = 3.07, p = 0.02, **Fig. 2a**). Shooting error (distance between the center of the target and the center of the sight) was 4.37 ± 1.19 minutes of arc in the normal condition, whereas it was 6.51 ± 1.09 minutes of arc in the eccentric condition (paired t-test, t(6) = −12.99, p < 0.01, **Fig. 2b**).

**Figure 2.**
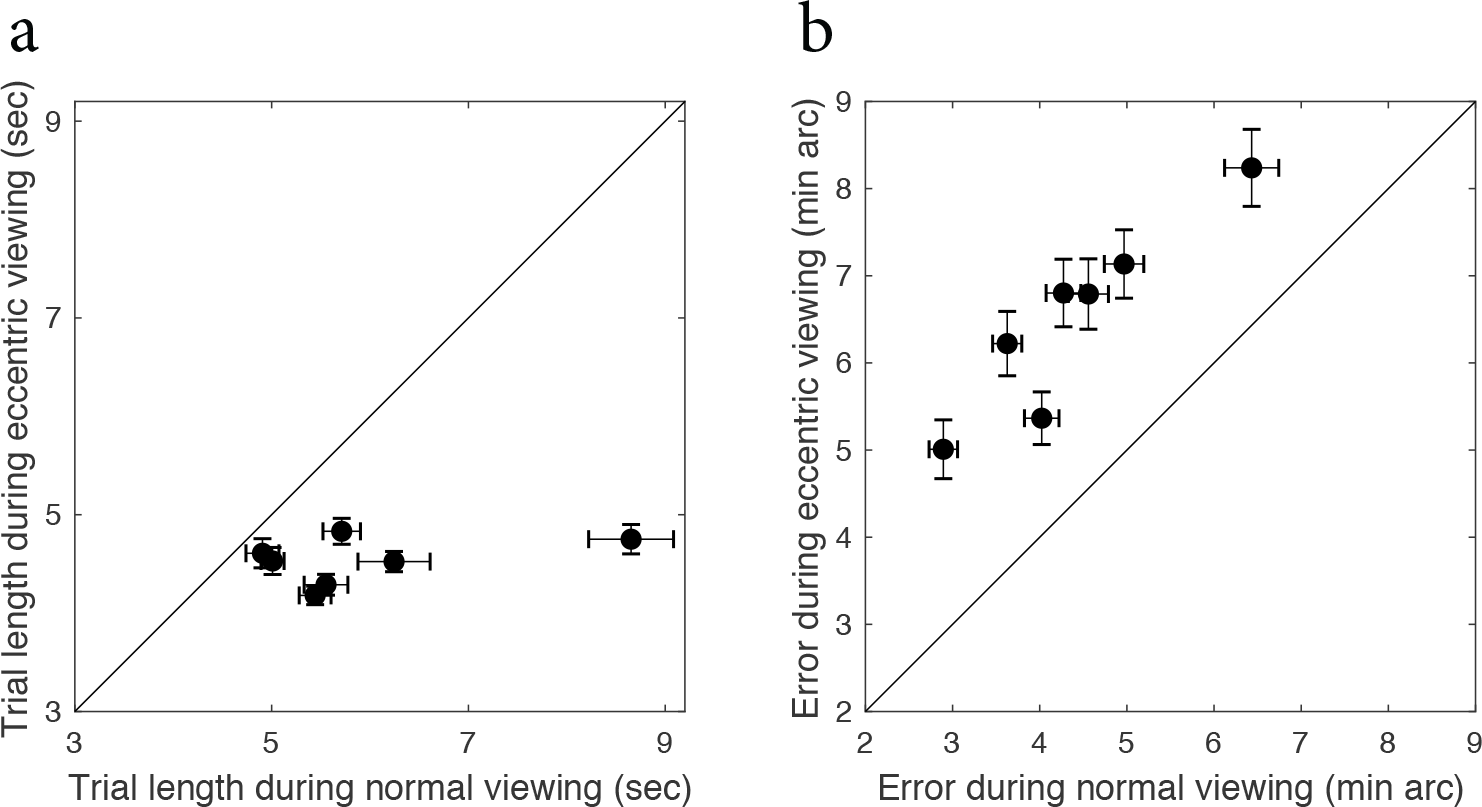
Time taken to shoot, and performance under normal and eccentric viewing conditions. (a) Each data point shows the mean time taken by a single participant to complete a trial under the two viewing conditions (n=7). Error bars represent s.e.m. (b) Each point shows for a single participant the mean error in alignment of the centers of the shooting target and sight when subjects pressed the “shoot” button, under the two viewing conditions.

The distribution of microsaccade amplitudes was also affected by task condition (Kruskal-Wallis test, H = 1660.72, df = 2, p < 0.01; **Fig. 3a**): specifically, participants made larger microsaccades when they performed the shooting task compared to when they just fixated on a fixation target (Tukey’s HSD, p < 0.01). On the other hand, there was no significant difference between the distribution of microsaccade amplitudes under the two viewing conditions of shooting (p = 0.2). To verify whether microsaccades were used to shift gaze between the two ROIs during the shooting task, we compared microsaccade amplitude as a function of the separation between the ROIs at beginning of each microsaccade. In the normal viewing condition, microsaccade amplitude was strongly correlated with ROI separation (**Fig. 3b**; r = 0.87, p < 0.01), which suggests that participants calibrated their microsaccade length to shift gaze between the ROIs. In the eccentric viewing condition, it is not possible to align gaze with the target or the sight, and, correspondingly, microsaccade amplitude was not correlated with the separation between ROIs (r = −0.36, p = 0.27).

**Figure 3.**
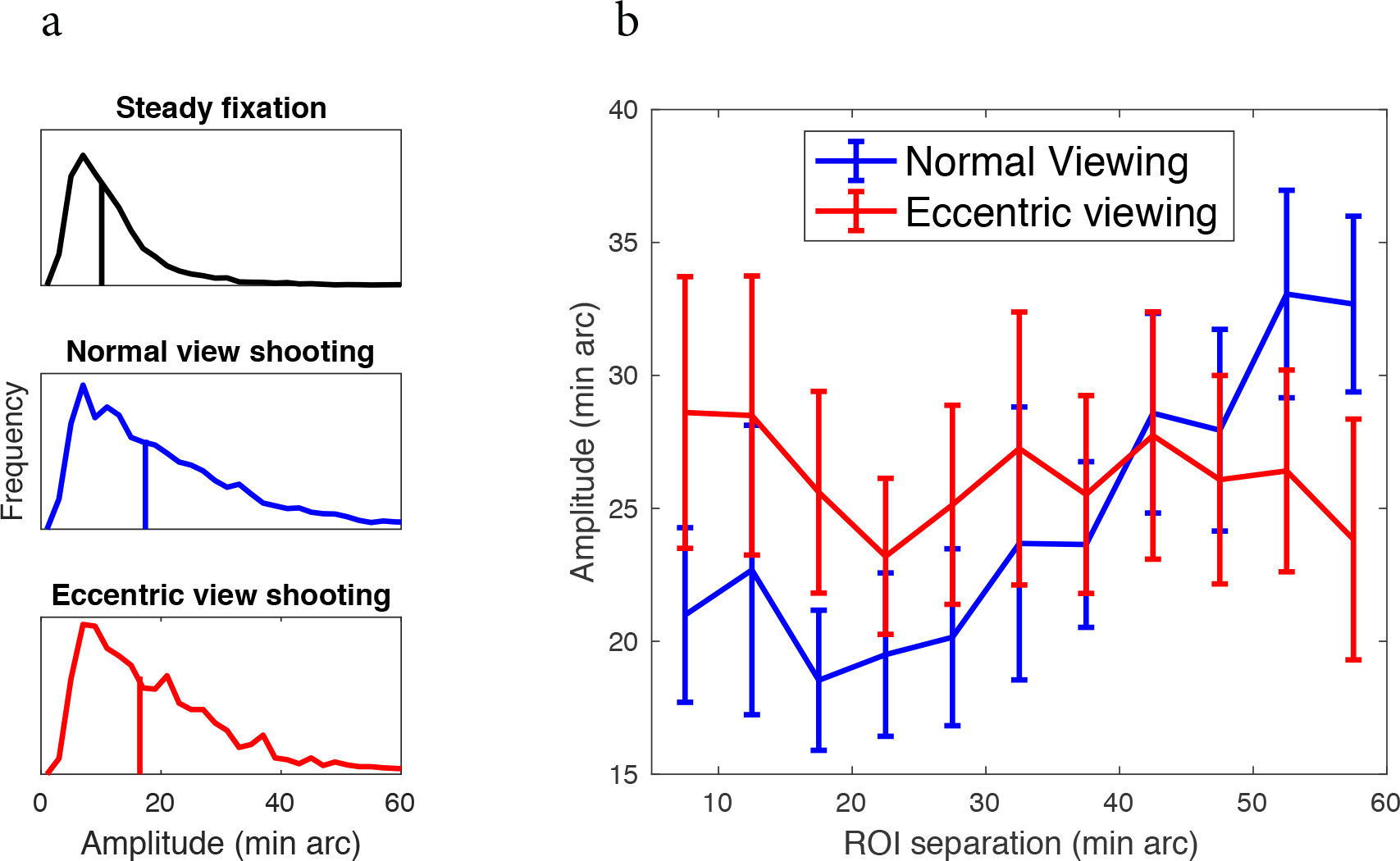
Comparison of microsaccade amplitude. (a) Microsaccade amplitude distribution for the different task conditions. The vertical lines show the median amplitude for each condition. (b) Microsaccade amplitude as a function of the separation between regions of interest at the time of execution of microsaccade. Error bars represent s.e.m.

We were interested to see whether there existed any temporal relationship between microsaccade occurrence and sight adjustment (through button press) events. For a given sight adjustment within a given trial, we subtracted all microsaccade starting times for that trial from the sight adjustment time and chose the one with the minimum absolute value as the microsaccade nearest to this sight adjustment in time. We did this for all sight adjustments within each viewing condition and obtained the corresponding distribution of nearest microsaccade times in relation to all sight adjustments (blue and red solid traces in **Fig. 4b**). A positive value means that a microsaccade followed the sight adjustment in time, while a negative value means that the microsaccade preceded the sight adjustment.

**Figure 4.**
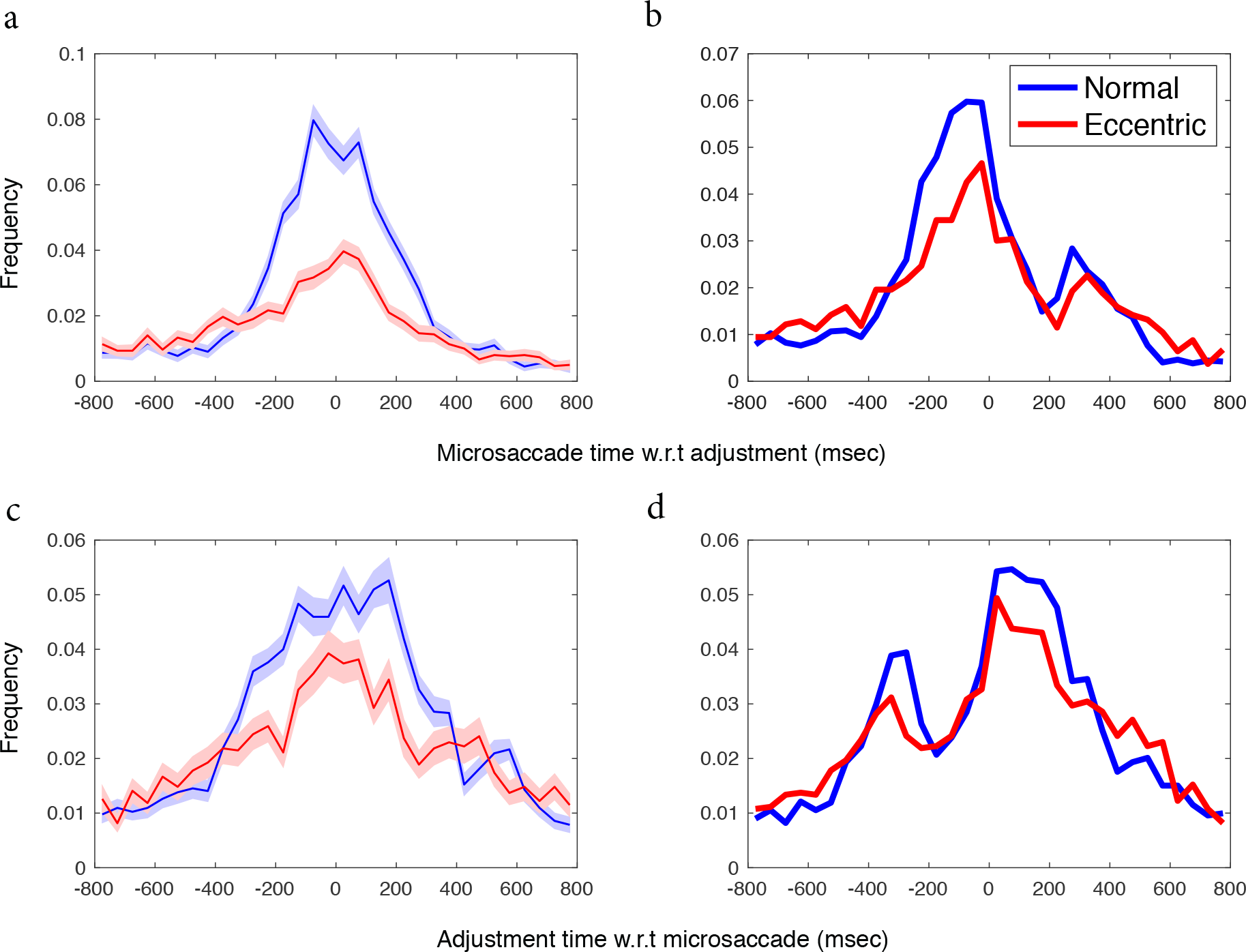
Temporal relation between microsaccade occurrence and sight adjustment times (button presses). (a) Solid lines show the mean of distribution of nearest pseudo-microsaccade times w.r.t each adjustment. Shaded area represents s.e.m. Blue: Normal viewing condition, Red: Eccentric viewing condition. (b) Distribution of nearest real-microsaccade times w.r.t to each adjustment. (c) Solid lines show the mean of distribution of nearest pseudo-button-press time w.r.t to each microsaccade, shaded region shows s.e.m. (d) Distribution of nearest real button-press times w.r.t each microsaccade.

For comparison, we repeated the above calculation with pseudo-microsaccades (**Fig. 4a**): a set of random time points across the duration of each trial. The number of such random points for a trial was same as the number of microsaccades in that trial. We repeated this process 100 times to get 100 frequency distributions of nearest pseudo-microsaccade times with respect to sight adjustment times. The probability of a microsaccade preceding an adjustment was higher in the time period ∼200 msec before an adjustment, whereas the probability of a microsaccade following an adjustment dropped during ∼200ms time period following an adjustment. This observation was true for both conditions. Similarly, we created distribution of adjustment time nearest to each microsaccade by subtracting each sight adjustment time from the microsaccade start time and picking the one with minimum absolute value as the sight adjustment nearest to this microsaccade in time (**Fig. 4d**). We compared this distribution with the distribution of pseudo-adjustment times computed as mentioned above for pseudo-microsaccade times (**Fig. 4c**). For both conditions, the probability of an adjustment dropped in the time period ∼200 msec preceding a microsaccade, whereas the probability of an adjustment increased in the time period ∼200 msec following a microsaccade. The two findings suggest that irrespective of the different viewing conditions, microsaccade and sight adjustment through button presses were temporally related, with adjustments remaining suppressed during the 200 msec time preceding a microsaccade, and occurring with a higher probability in the ∼200 msec following a microsaccade.

Next, we compared mean microsaccade rate near the end of the shooting task for the two viewing conditions by aligning each trial’s microsaccade rate function to trial end (**Fig 5a**). In the normal viewing condition, mean microsaccade rate in the final second of the task was significantly lower than in the third second before trial end (one tailed paired t-test, t (6) = −3.34, p < 0.01, **Fig. 5b**). This drop in microsaccade rate near the end of the shooting task agrees with the results of previous studies (e.g., Bridgeman & Palca, 1980; Ko, Poletti, & Rucci, 2010; Winterson & Collewun, 1976), and has been attributed to the lack of a need to shift gaze as the ROIs start to overlap on the effective foveal region (Ko et al., 2010). Surprisingly, mean microsaccade rate also decreased toward the end of the trial during the eccentric viewing condition (t(6) = −4.91, p < 0.01), even though in this task, microsaccades could not function to shift gaze between the relevant task objects. This indicates that in the shooting task, microsaccade rate is modulated toward the end of the trial in a similar way, irrespective of whether they performed a gaze orientating function (as in the normal viewing condition) or not (as in the eccentric viewing condition). For comparison, we also analyzed microsaccade rate during the fixation task. Since the end of the 5-second fixation period was followed by a shooting task trial, we selected a 3-second time period from the middle of each fixation trial for analysis. This avoided possible modulations in microsaccade rate due to the anticipated onset of the subsequent trial. Microsaccade rate during fixation period remained unchanged (t(6) = −0.48, p = 0.32).

**Figure 5.**
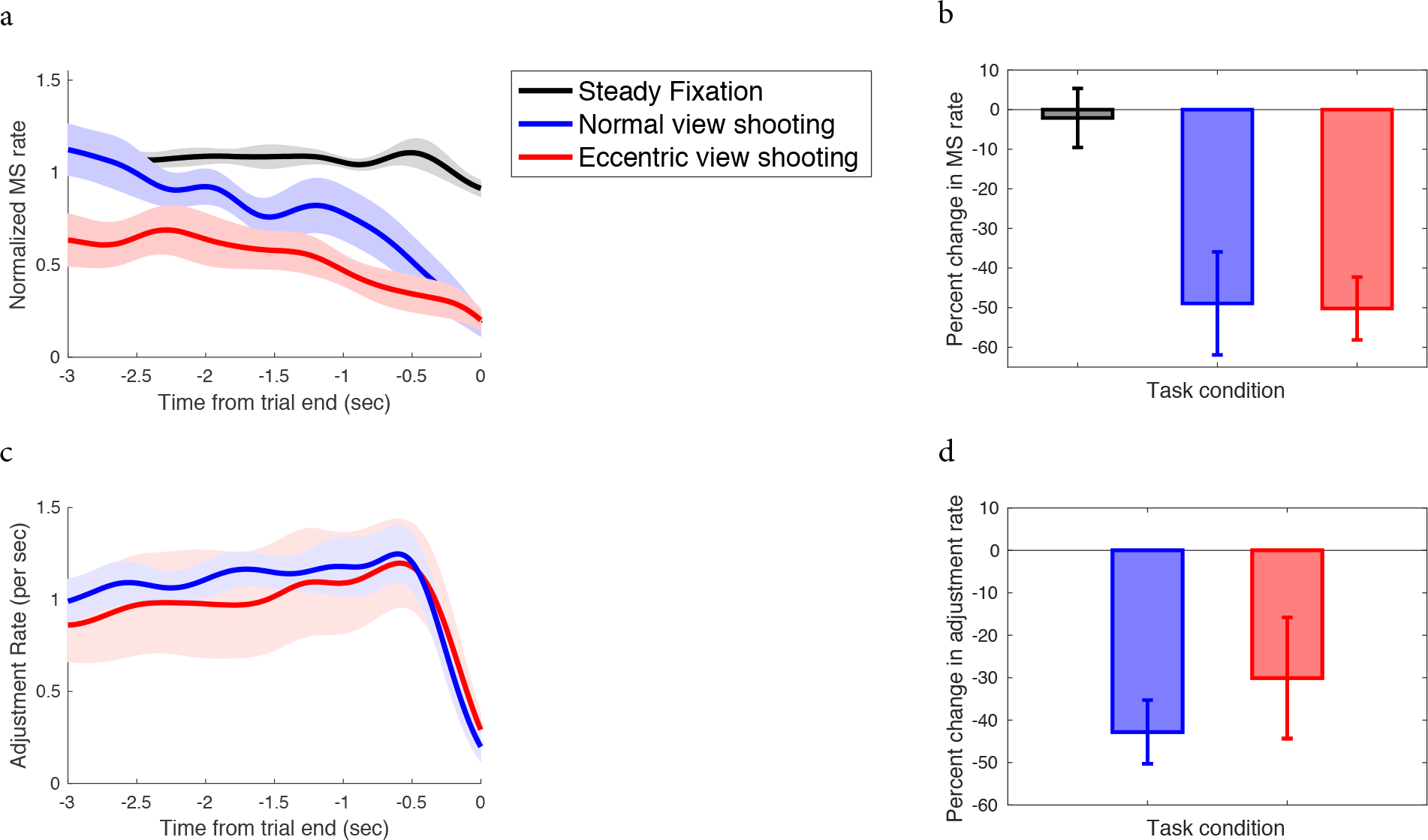
Mean microsaccade and adjustment (button press) rate aligned to trial end. (a) Solid lines show average of all participants’ mean normalized microsaccade rate during steady fixation (black), shooting under normal viewing (blue) and eccentric viewing (red), aligned to trial end. Shaded region indicates s.e.m. (b) Percent change in microsaccade rate from early period (−3 to −2 sec from trial end) to the end period (last 1 sec). Error bars represent s.e.m. Color coding of bars is same as in panel (a). (c) Rate of button presses resulting in adjustment of sight’s direction; aligned to trial end. (d) Percent change in adjustment rate from an early time (−1 to −0.5 sec from trial end) to trial end (last 0.5 sec). Color coding in (b), (c), and (d) same as in (a).

If the earlier reported and currently observed drop in microsaccade rate at the end of shooting task is because of the two ROIs finally overlapping on the same effective foveal region and thus making the need to shift gaze obsolete or even harmful for the task, then what could have caused the drop in microsaccade rate in the eccentric viewing condition, when microsaccades did not occur to overtly shift gaze between the relevant task objects? One possibility is that this drop in microsaccade rate may be related to a change in the allocation of covert spatial attention near the end of the trial.

Microsaccades have been shown to reflect shifts in covert attention (Engbert & Kliegl, 2003b). Immediately following a covert shift of attention, microsaccade probability increases and their directions tend to align with the direction of the newly attended location. Based on this, one possible explanation for the observed drop in microsaccade rate during eccentric viewing condition could be this: during the initial phase of the trial, participants constantly switch attention alternatively between the central fixation target and eccentric task location, and the microsaccades thus produced are reflective of this phenomenon. Nearing the end of trial, at a certain time before actually pressing the shoot button, participants may stop assessing the alignment between the shooting stimuli as the poor quality of peripheral vision does not afford fine judgement of the alignment of the two objects. Thus, participants pre-decide when to shoot and disengage attention from the eccentric task location before the end of the task, and this is reflected by the decrease in microsaccadic activity at the end. We conducted the next experiment to test this hypothesis.

In a dual attention task, participants detected a peripheral target letter from a rapid stream of letters (RSVP task; Rapid Serial Visual Presentation), while simultaneously performing the same shooting task as before. They viewed the shooting stimuli under the same two conditions as earlier; normal and eccentric. The RSVP stimuli appeared at an eccentric location in both conditions, and, for the eccentric viewing condition, this location was always in the hemifield opposite the shooting stimuli. If the attentional disengagement hypothesis is true, then in the eccentric viewing condition, it would be expected that performance in the RSVP task would improve at end of the trial as attention is disengaged from shooting task and is readily available to be allocated to RSVP task location.

Microsaccade rates showed a similar pattern as during the first experiment, with their rate decreasing significantly in the later part of the trial for both the normal and eccentric viewing conditions (**Fig 6a**). However, RSVP task performance did not improve toward the end of the trial in the eccentric viewing condition (**Fig 6b**). Instead, it deteriorated with time until the end of the trial for both conditions. This suggests that attentional disengagement from the shooting target location is not the explanation for the observed drop in microsaccade rate at the end of the trial in the eccentric viewing condition.

**Figure 6.**
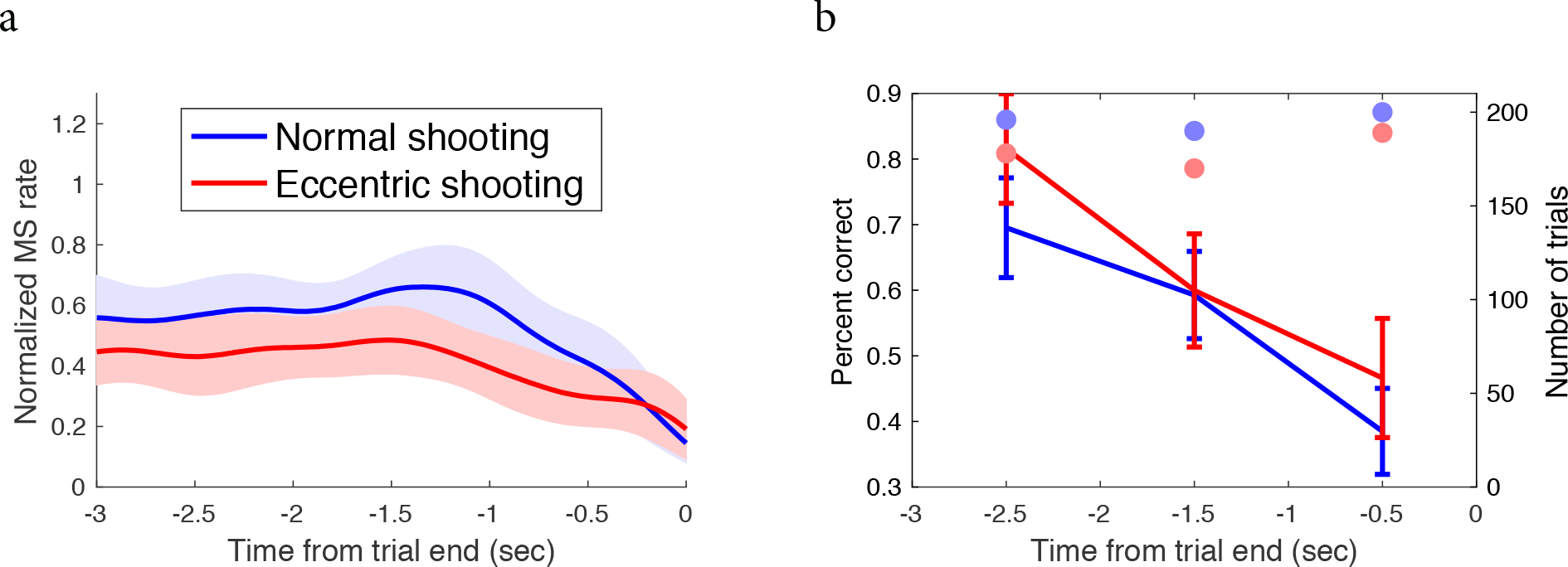
RSVP Performance in dual task. (a) Mean normalized microsaccade rate aligned to trial end for shooting under normal viewing (blue) and eccentric viewing (red) in the dual task. RSVP task was at an eccentric location in both conditions, and opposite to shooting location in the eccentric shooting condition. Shaded regions represent s.e.m. (b) Target letter detection in RSVP task under the two viewing conditions of shooting in dual task. Color coding same as in (a). Error bars represent s.e.m. Colored dots in each time bin indicate the total number of trials in which RSVP target appeared at corresponding times. Blue; normal shooting condition, red; eccentric shooting condition.

## DISCUSSION

While performing tasks requiring fine acuity like shooting or threading a needle, microsaccades initially shift gaze precisely between the task relevant objects over a very narrow range of the visual field (< 1°; Ko, Poletti, & Rucci, 2010), just like bigger saccades (Otero-Millan, Macknik, Langston, & Martinez-Conde, 2013), and during the final second of the task their rate drops drastically, with no microsaccades at all in most of the trials (Winterson & Collewun, 1976; Bridgeman & Palca, 1980; Ko et al., 2010). This drop in microsaccade rate within a trial, along with the lower average rate during a task, as compared to during fixation, had initially been explained as a voluntary suppression of microsaccades to avoid their effects on visual perception, which could deteriorate performance (Winterson & Collewun, 1976). But a more recent study (Ko et al., 2010) suggested that as the relevant task objects start to overlap in the foveal region, the need to shift gaze is obviated, resulting in a drop in their rate.

Here, we asked human participants to perform a simulated shooting task in a normal viewing condition, in which they were free to move their eyes, and in an eccentric viewing condition, in which they fixated on a central target while performing the same task at an eccentric location. The two different conditions dissociated the gaze-reorienting function of microsaccades, as microsaccades made at the fixation location in the eccentric condition could be spontaneous or a result of sustained covert attention (Pastukhov, Vonau, Stonkute, & Braun, 2013), but could not reorient gaze between the shooting task stimuli. Thus, any modulation in microsaccade rate in this condition would be independent of its gaze-orienting function. For the normal shooting condition, we replicated the findings of earlier studies. Surprisingly, even for the eccentric condition, microsaccade rate showed a fairly abrupt drop during the final second of the task (Fig. 5), indicating that subjects suppressed microsaccades even when they did not shift gaze between the task relevant objects. Although the drop in microsaccade rate in the normal viewing condition can be explained by the demand of dynamic gaze shifts fulfilled by microsaccades (Ko et al., 2010), it does not explain why such a drop would be observed in the eccentric viewing condition.

Possible explanations for the observed drop in microsaccade rate in our study are: spatial disengagement of covert attention from eccentric task location, active suppression to improve task performance, preparation or execution of oculomotor (saccade towards eccentric task location to judge task performance after the task ends) or manual (key press to ‘shoot’) response, perceptual decision making process, temporal expectation, and changes in temporal attention. We will go through these possibilities one by one.

Participants performed the shooting task less accurately in the eccentric condition (Fig. 2), consistent with poorer localization of the task stimuli when viewed in peripheral vision. Hence it is possible that they focused more on the initial coarser adjustments of the gun sight and then pressed the trigger at a pre-planned time based on the sight’s velocity information, without making final fine adjustments. Since participants in this condition fixated a central target while covertly paying attention to the eccentric task location, microsaccades produced initially in a trial could have been the result of alternating attention between the fixation target and eccentric task location, as it has been shown by both psychophysical (Engbert & Kliegl, 2003a; Hafed & Clark, 2002; Hafed, Lovejoy, & Krauzlis, 2011; Pastukhov et al., 2013) and electrophysiological (Hafed, Lovejoy, & Krauzlis, 2013) studies that microsaccade characteristics are related to immediate and sustained shifts of covert attention. The observed drop in microsaccade rate in the eccentric viewing condition could have resulted from a disengagement of attention from the peripheral location toward the end of the task. To test this possibility, we asked subjects to perform the shooting task under the same two conditions, but with an added RSVP task at an eccentric and opposite location. If attention was indeed disengaged from the eccentric shooting task location, then in this dual task situation, we expected greater attention resources would be available to be captured by the RSVP task location, thus making performance in the RSVP task better when microsaccade rate decreased in the peripheral task. On the other hand, in the normal condition we expected no such change in RSVP performance. We found that RSVP performance near the end of the trial deteriorated in both the conditions (Fig. 6), suggesting that an attentional disengagement from the shooting stimuli is unlikely to be the explanation for the observed drop in microsaccade rate. Nonetheless, it is also possible that our RSVP task failed to capture the disengaged attention or that attention in that short span was, instead, diffused over the entire field, or that attention was instead focused primarily on the central fixation target. In all such cases microsaccade rate would still be expected to drop. Carefully designed experiments would be needed in the future to test each of these specific hypotheses.

Our results show that microsaccades are suppressed during the end of the shooting task, just before fine acuity information is presumably processed to judge the relative alignment of the target and sight to arrive at a perceptual decision as to when to press the trigger. Like larger saccades, microsaccades also modulate visual perception around the time of their occurrence, which could affect task performance adversely (Hafed, Chen, & Tian, 2015). They compress the perception of both space (Hafed, 2013) and time (Yu, Yang, Yu, & Dorris, 2017). To avoid the adverse effects of such perceptual modulations on task performance, the oculomotor system might have learnt to suppress microsaccades at a time when a perceptual decision has to be made (Bridgeman & Palca, 1980). Hafed et al. (2011) found that on a sustained covert attention task requiring monkeys to judge the direction of a motion pulse at a cued location, monkeys took longer to respond and tended to make more errors if a microsaccade occurred near the time of motion pulse onset. Similarly, Xue et al. (2017) more recently found that in a color change detection task, microsaccades occurring in the period ±100 ms around the color change delayed the monkey’s response time and also worsened their detection performance. A lower microsaccade rate is also correlated with higher accuracy in orientation judgements (Amit, Abeles, Carrasco, & Yuval-Greenberg, 2019; Denison, Yuval-Greenberg, & Carrasco, 2019). We did not find any correlation between the timing of the last microsaccade and performance in a trial. Our failure to find any relation between microsaccade occurrence and task performance could be because of three reasons: first, as opposed to monkeys, humans participated in our experiment, and it is possible that humans, through learning, have become more robust in performance in such tasks. Second, instead of detection, our task required humans to constantly judge the relative positions of the task objects to finally make a perceptual judgement about when the two objects were concentric, and then press a trigger. Microsaccades might have impaired the perceptual view of a stimulus change around their time of occurrence when the response is binary (yes/no), but they might not affect performance in our task in which performance in judged on a continuous scale (by how much distance was the target missed?). Finally, even if the occurrence of microsaccades did affect performance, our experimental design might have failed to capture it because the number of trials in which microsaccades did occur near the end was smaller than the number of trials in which there were no microsaccades, and thus, the margin of error may have been too small to observe any significant difference in performance.

Microsaccades are suppressed in preparation of saccades (Rolfs, Laubrock, & Kliegl, 2006; Hermens, Zanker, & Walker, 2010; Watanabe, Matsuo, Zha, Munoz, & Kobayashi, 2013; Dalmaso, Castelli, & Galfano, 2019) as well as manual responses (Betta & Turatto, 2006). In our present study, participants did not make a saccade towards the eccentric task objects since we enforced fixation on the fixation target using an invisible fixation boundary. Also, the shooting score in any given trial was presented at the fixation target location at the end of the trial, thus obviating the need to make a saccade towards am eccentric location. Any reflexive saccades made to the eccentric location terminated the trial even if the participant had already pressed the ‘shoot’ button, and the number of such trials was very low. Hence our analysis did not include such trials, and we can rule out the possibility of microsaccades getting suppressed by saccade preparation. Microsaccades have been shown to be suppressed during needle threading tasks even in the absence of manual response ((Bridgeman & Palca, 1980; Ko et al., 2010)). In these studies, participants passively watch a thread approaching the needle until the partial completion of the trial after which the task stimulus was masked and participants responded orally about their judgement about whether the thread made it through the needle’s eye or not. This allows us to rule out the possibility of a manual response preparation induced suppression of microsaccades.

The only explanations we are left out with are temporal expectation of a visual change and changes in temporal attention. Amit et al., (2019) showed that microsaccades are inhibited when a temporal cue is informational about an upcoming target whose orientation has to be judges. In addition the cue-target interval was also related to the intensity of microsaccade inhibition, reflecting a microsaccade suppression mechanism driven by anticipation of an event. This could well be the reason for the suppression of microsaccades in our task since a the end of every trial, participants anticipated their performance which presented to them in the form of a shooting score. Although we ruled out the possibility of change in the focus of spatial attention during the end of the task affecting microsaccade rate, it is possible that a change in temporal attention might have contributed to the drop in microsaccade rate. Such suppression has been reported recently in a an orientation discrimination task (Denison et al., 2019) in which a temporal cue is informative about which target among a stream of targets is to be assessed. It is possible that participants in our study directed their temporal attention more intensely nearing the end of the task in an attempt to judge the relative position of the target and sight when they were the nearest to each other which could have caused the suppression of microsaccades.

An interesting finding in our study is the tight temporal coupling between the microsaccades and button presses used to adjust sight direction (Fig. 4). Button presses were more likely to be executed in the ∼200 ms following a microsaccade, whereas microsaccades were inhibited in the ∼200 ms following an adjustment. Two inferences can be drawn from this observation; first, adjustments were mostly made after a microsaccade was made to judge the relative position of the target and sight, similar to Ko et al.'s (2010) findings. This provides evidence to support the idea that microsaccades serve a gaze-shifting function similar to larger saccades, which is then used to aid manual adjustments of the sight’s direction. Second, microsaccades and button presses always occurred in a particular sequence and not simultaneously, which suggests that oculomotor and motor (hand) response preparation could share a common cognitive resource (Betta & Turatto, 2006).

There are two limitations to our current study. First, we used a video-based eye tracker that has a relatively low spatial resolution, significant trial-to-trial variability in its estimate of actual line of sight (Kimmel, Mammo, & Newsome, 2012), and pupil size change induced artifacts in eye position signal (Nyström, Hooge, & Andersson, 2016). These factors limited our analysis of microsaccade directions. Thus, we cannot say with confidence the precise position of the actual line of sight in the range of the fine resolution of our task stimuli, or the exact task object to which the microsaccades were directed. Nonetheless our finding of a positive and significant correlation between the microsaccade amplitude and separation between the task objects only in the normal viewing condition is sufficient to infer that microsaccades were used to shift gaze precisely when they were free to move their eyes. Second, we assume that microsaccades made during the eccentric shooting condition do not contribute to the perception of object positions in the peripheral field of vision .. Hennig & Wörgötter (2003) through a model of the vertebrate retinal response to resting and moving eye suggested that eye movements in the range of microsaccade can contribute to peripheral acuity by reducing the effects of neural undersampling induced aliasing. Chen, Ignashchenkova, Thier, & Hafed (2015) showed a neural response gain enhancement for peripheral locations in superior colliculus and frontal eye field prior to the occurrence of microsaccades. Although these studies suggest that microsaccades might enhance visual processing at peripheral locations, there is no strong evidence to believe that subjects in our task could be using it to their advantage. We believe that microsaccades occurring during the eccentric viewing condition could just be spontaneous events, or indicators of sustained covert attention, or a mixture of both. In any case, enhancements in peripheral visual processing afforded by microsaccades, if any, would be very low compared to their contribution in the fine visual discrimination at the fovea.

Finally, we emphasize the developments made in the study of microsaccade’s role or the lack of it in various task scenarios. Initially the volitional control of microsaccade was suggestive of its role in oculomotor strategies (Steinman, Cunitz, Timberlake, & Herman, 1967; Winterson & Collewun, 1976; Steinman, Haddad, Skavenski, & Wyman, 1973) but Kowler & Steinman (1979) denied its role in cognitive tasks like counting. Most of the later studies focused on microsaccadic response to low level processes like transient visual display changes or exogenous attention shifts (Hafed & Clark, 2002; Engbert & Kliegl, 2003a). Recent studies have tried to expand the role of microsaccades in more complex tasks that require higher level functions. Betta & Turatto (2006) showed that microsaccade rate is linked to manual response preparation and can be a measure of readiness. Valsecchi, Betta, & Turatto (2007) showed that the characteristic microsaccade rate signature is temporally expanded in a visual odd ball task and the effect is more pronounced when the oddball is task relevant. Hafed et al. (2011) showed through a monkey study that microsaccades are aligned to the axis of sustained covert attention and might be used by the visual and oculomotor system as part of a sophisticated fixation strategy. Microsaccade studies have also expanded to the field of object categorization (Craddock, Oppermann, Müller, & Martinovic, 2017), overt attentional selection (Meyberg, Sinn, Engbert, & Sommer, 2017), perceived compression of time (Yu et al., 2017), working memory (Dalmaso, Castelli, Scatturin, & Galfano, 2017), spatiotemporal information processing (Boi, Poletti, Victor, & Rucci, 2017), reading (Bowers & Poletti, 2017; Yablonski, Polat, Bonneh, & Ben-Shachar, 2017), idea generation (Walcher, Körner, & Benedek, 2017) and music absorption (Lange, Zweck, & Sinn, 2017). In the light of these recent studies, our current study adds to the evidence that microsaccade characteristics not only reflect low level processing, but they are also modulated during tasks involving higher cognitive processing.

## CONCLUSION

Microsaccades are suppressed during the execution of fine acuity tasks like shooting even when they do not contribute in shifting gaze over task relevant objects. Such suppression might be reflective of the cognitive processes involved in such tasks like perceptual decision making and response preparation.

## REFERENCES

Amit, R., Abeles, D., Carrasco, M., & Yuval-Greenberg, S. (2019). Oculomotor inhibition reflects temporal expectations. NeuroImage, 184, 279–292. https://doi.org/10.1016/j.neuroimage.2018.09.026

Betta, E., & Turatto, M. (2006). Are you ready? I can tell by looking at your microsaccades. NeuroReport, 17(10), 1001. https://doi.org/10.1097/01.wnr.0000223392.82198.6d

Boi, M., Poletti, M., Victor, J. D., & Rucci, M. (2017). Consequences of the Oculomotor Cycle for the Dynamics of Perception. Current Biology, 27(9), 1268–1277. https://doi.org/10.1016/j.cub.2017.03.034

Bowers, N. R., & Poletti, M. (2017). Microsaccades during reading. PLOS ONE, 12(9), e0185180. https://doi.org/10.1371/journal.pone.0185180

Bridgeman, B., & Palca, J. (1980). The role of microsaccades in high acuity observational tasks. Vision Research, 20(9), 813–817. https://doi.org/10.1016/0042-6989(80)90013-9

Chen, C.-Y., Ignashchenkova, A., Thier, P., & Hafed, Z. M. (2015). Neuronal Response Gain Enhancement prior to Microsaccades. Current Biology, 25(16), 2065–2074. https://doi.org/10.1016/j.cub.2015.06.022

Collewijn, H., & Kowler, E. (2008). The significance of microsaccades for vision and oculomotor control. Journal of Vision, 8(14), 20–20. https://doi.org/10.1167/8.14.20

Craddock, M., Oppermann, F., Müller, M. M., & Martinovic, J. (2017). Modulation of microsaccades by spatial frequency during object categorization. Vision Research, 130, 48–56. https://doi.org/10.1016/j.visres.2016.10.011

Dalmaso, M., Castelli, L., & Galfano, G. (2019). Microsaccadic rate and pupil size dynamics in pro-/anti-saccade preparation: the impact of intermixed vs. blocked trial administration. Psychological Research. https://doi.org/10.1007/s00426-018-01141-7

Dalmaso, M., Castelli, L., Scatturin, P., & Galfano, G. (2017). Working memory load modulates microsaccadic rate. Journal of Vision, 17(3), 6–6. https://doi.org/10.1167/17.3.6

Denison, R. N., Yuval-Greenberg, S., & Carrasco, M. (2019). Directing Voluntary Temporal Attention Increases Fixational Stability. Journal of Neuroscience, 39(2), 353–363. https://doi.org/10.1523/JNEUROSCI.1926-18.2018

Dorr, M., Martinetz, T., Gegenfurtner, K. R., & Barth, E. (2010). Variability of eye movements when viewing dynamic natural scenes. Journal of Vision, 10(10), 28–28. https://doi.org/10.1167/10.10.28

Engbert, R., & Kliegl, R. (2003a). Microsaccades uncover the orientation of covert attention. Vision Research, 43(9), 1035–1045. https://doi.org/10.1016/S0042-6989(03)00084-1

Engbert, R., & Kliegl, R. (2003b). Microsaccades uncover the orientation of covert attention. Vision Research, 43(9), 1035–1045. https://doi.org/10.1016/S0042-6989(03)00084-1

Hafed, Z. M. (2013). Alteration of Visual Perception prior to Microsaccades. Neuron, 77(4), 775–786. https://doi.org/10.1016/j.neuron.2012.12.014

Hafed, Z. M., Chen, C.-Y., & Tian, X. (2015). Vision, Perception, and Attention through the Lens of Microsaccades: Mechanisms and Implications. Frontiers in Systems Neuroscience, 9. https://doi.org/10.3389/fnsys.2015.00167

Hafed, Z. M., & Clark, J. J. (2002). Microsaccades as an overt measure of covert attention shifts. Vision Research, 42(22), 2533–2545. https://doi.org/10.1016/S0042-6989(02)00263-8

Hafed, Z. M., Lovejoy, L. P., & Krauzlis, R. J. (2011). Modulation of Microsaccades in Monkey during a Covert Visual Attention Task. Journal of Neuroscience, 31(43), 15219–15230. https://doi.org/10.1523/JNEUROSCI.3106-11.2011

Hafed, Z. M., Lovejoy, L. P., & Krauzlis, R. J. (2013). Superior colliculus inactivation alters the relationship between covert visual attention and microsaccades. The European Journal of Neuroscience, 37(7), 1169–1181. https://doi.org/10.1111/ejn.12127

Hennig, M. H., & Wörgötter, F. (2003). Eye Micro-Movements Improve Stimulus Detection beyond the Nyquist Limit in the Peripheral Retina.

Hermens, F., Zanker, J. M., & Walker, R. (2010). Microsaccades and preparatory set: a comparison between delayed and immediate, exogenous and endogenous pro- and anti-saccades. Experimental Brain Research, 201(3), 489–498. https://doi.org/10.1007/s00221-009-2061-5

Kimmel, D. L., Mammo, D., & Newsome, W. T. (2012). Tracking the eye non-invasively: simultaneous comparison of the scleral search coil and optical tracking techniques in the macaque monkey. Frontiers in Behavioral Neuroscience, 6. https://doi.org/10.3389/fnbeh.2012.00049

Ko, H., Poletti, M., & Rucci, M. (2010). Microsaccades precisely relocate gaze in a high visual acuity task. Nature Neuroscience, 13(12), 1549–1553. https://doi.org/10.1038/nn.2663

Kowler, E., & Steinman, R. M. (1979). Miniature saccades: eye movements that do not count. Vision Research, 19(1), 105–108.

Lange, E. B., Zweck, F., & Sinn, P. (2017). Microsaccade-rate indicates absorption by music listening. Consciousness and Cognition, 55, 59–78. https://doi.org/10.1016/j.concog.2017.07.009

Meyberg, S., Sinn, P., Engbert, R., & Sommer, W. (2017). Revising the link between microsaccades and the spatial cueing of voluntary attention. Vision Research, 133, 47–60. https://doi.org/10.1016/j.visres.2017.01.001

Nyström, M., Hooge, I., & Andersson, R. (2016). Pupil size influences the eye-tracker signal during saccades. Vision Research, 121, 95–103. https://doi.org/10.1016/j.visres.2016.01.009

Otero-Millan, J., Macknik, S. L., Langston, R. E., & Martinez-Conde, S. (2013). An oculomotor continuum from exploration to fixation. Proceedings of the National Academy of Sciences, 110(15), 6175–6180. https://doi.org/10.1073/pnas.1222715110

Pastukhov, A., Vonau, V., Stonkute, S., & Braun, J. (2013). Spatial and temporal attention revealed by microsaccades. Vision Research, 85, 45–57. https://doi.org/10.1016/j.visres.2012.11.004

Rolfs, M. (2009). Microsaccades: Small steps on a long way. Vision Research, 49(20), 2415–2441. https://doi.org/10.1016/j.visres.2009.08.010

Rolfs, M., Laubrock, J., & Kliegl, R. (2006). Shortening and prolongation of saccade latencies following microsaccades. Experimental Brain Research, 169(3), 369–376. https://doi.org/10.1007/s00221-005-0148-1

Steinman, R. M., Cunitz, R. J., Timberlake, G. T., & Herman, M. (1967). Voluntary Control of Microsaccades during Maintained Monocular Fixation. Science, 155(3769), 1577–1579. https://doi.org/10.1126/science.155.3769.1577

Steinman, R. M., Haddad, G. M., Skavenski, A. A., & Wyman, D. (1973). Miniature Eye Movement. Science, 181(4102), 810–819. https://doi.org/10.1126/science.181.4102.810

Valsecchi, M., Betta, E., & Turatto, M. (2007). Visual oddballs induce prolonged microsaccadic inhibition. Experimental Brain Research, 177(2), 196–208. https://doi.org/10.1007/s00221-006-0665-6

Walcher, S., Körner, C., & Benedek, M. (2017). Looking for ideas: Eye behavior during goal-directed internally focused cognition. Consciousness and Cognition, 53, 165–175. https://doi.org/10.1016/j.concog.2017.06.009

Watanabe, M., Matsuo, Y., Zha, L., Munoz, D. P., & Kobayashi, Y. (2013). Fixational saccades reflect volitional action preparation. Journal of Neurophysiology, 110(2), 522–535. https://doi.org/10.1152/jn.01096.2012

Winterson, B. J., & Collewun, H. (1976). Microsaccades during finely guided visuomotor tasks. Vision Research, 16(12), 1387–1390. https://doi.org/10.1016/0042-6989(76)90156-5

Xue, L., Huang, D., Wang, T., Hu, Q., Chai, X., Li, L., & Chen, Y. (2017). Dynamic modulation of the perceptual load on microsaccades during a selective spatial attention task. Scientific Reports, 7(1), 16496. https://doi.org/10.1038/s41598-017-16629-2

Yablonski, M., Polat, U., Bonneh, Y. S., & Ben-Shachar, M. (2017). Microsaccades are sensitive to word structure: A novel approach to study language processing. Scientific Reports, 7(1), 3999. https://doi.org/10.1038/s41598-017-04391-4

Yu, G., Yang, M., Yu, P., & Dorris, M. C. (2017). Time compression of visual perception around microsaccades. Journal of Neurophysiology, 118(1), 416–424. https://doi.org/10.1152/jn.00029.2017

